# Cognitive and GABAergic protection by hAPP^Sw,Ind^ and hTau^VLW^ coexpression: a model of Non-Demented with Alzheimer’s disease Neuropathology

**DOI:** 10.1101/2020.12.10.417907

**Authors:** Eva Dávila-Bouziguet, Arnau Casòliba-Melich, Georgina Targa-Fabra, Lorena Galera-López, Andrés Ozaita, Rafael Maldonado, José M. Delgado-García, Agnès Gruart, Jesús Ávila, Eduardo Soriano, Marta Pascual

## Abstract

Alzheimer’s disease comprises amyloid-β (Aβ) and hyperphosphorylated Tau (P-Tau) accumulation, imbalanced neuronal activity, aberrant oscillatory rhythms, and cognitive deficits. Non-Demented with Alzheimer’s disease Neuropathology (NDAN) defines a novel clinical entity with Aβ and Tau pathologies, but preserved cognition. The mechanisms underlying such neuroprotection remain undetermined and animal models are currently unavailable for NDAN. We show that J20/VLW mice, accumulating Aβ and P-Tau, exhibit preserved hippocampal rhythmic activity and cognition, altered in J20 and VLW animals. Furthermore, we show that coexistence with Aβ renders a particular P-Tau signature in hippocampal interneurons. The GABAergic septohippocampal pathway, responsible for hippocampal rhythmic activity, is preserved in J20/VLW mice, in contrast to single mutants. Our data highlight J20/VLW mice as a suitable animal model to understand the mechanisms driving cognitive preservation in NDAN and suggest that a differential P-Tau pattern in hippocampal interneurons prevents GABAergic septohippocampal innervation loss and alterations in local field potentials, avoiding cognitive deficits.

## INTRODUCTION

The histopathological hallmarks of Alzheimer’s disease (AD) include neurodegeneration, extracellular deposits of amyloid-β peptide (Aβ; i.e. senile plaques), and intracellular neurofibrillary tangles of hyperphosphorylated Tau protein (P-Tau) (Kametani and Hasegawa, 2018). In addition, altered neural activity, such as synaptically driven hyperactivity, has been observed in mouse models of AD (Busche et al., 2012; Palop et al., 2007), and functional MRI studies have also revealed hippocampal hyperactivity in asymptomatic individuals at genetic risk of AD (Bookheimer et al., 2000; Reiman et al., 2012). Moreover, epileptiform activity is more common in individuals with AD than in controls (Horváth et al., 2016). Theta and gamma oscillations of local field potentials (LFPs) are reduced in AD patients and animal models, thereby further indicating an imbalance between excitatory and inhibitory circuits (Mably and Colgin, 2018; Palop and Mucke, 2016; Rubio et al., 2012; Verret et al., 2012). Theta and gamma oscillations are regulated by fast-spiking, Parvalbumin (PV)-positive hippocampal interneurons (Buzsáki, 2002; Sohal et al., 2009). At the circuit level, hippocampal PV-positive cells provide perisomatic inhibition onto glutamatergic pyramidal and granular neurons and regulate spike timing (Amilhon et al., 2015). Consistent with reduced PV output in AD, pyramidal neurons show reduced spontaneous GABAergic currents, and optogenetic stimulation of PV-positive cells improves gamma oscillations and reduces Aβ load and P-Tau levels in AD mouse models (Etter et al., 2019; Iaccarino et al., 2016). Additional work suggests that loss of GABAergic tone partly underlies network dysfunction in AD and tauopathies (Ambrad Giovannetti and Fuhrmann, 2019; Busche et al., 2015; Shimojo et al., 2020).

The medial septum and diagonal band of Broca (MSDB) complex and the nucleus basalis of Meynert innervate the cerebral cortex and the hippocampus. Along with cholinergic fibers, the septohippocampal (SH) pathway consists of another key element, namely the GABAergic SH pathway. This component involves inhibitory long-range projection neurons that terminate specifically on GABAergic hippocampal interneurons (Freund and Antal, 1988; Gulyás et al., 1990), which in turn govern the activity of pyramidal neurons. The activation of GABAergic SH neurons is believed to result in the selective inhibition of inhibitory interneurons, hence enabling the synchronous activation of a large number of pyramidal neurons (Freund and Gulyás, 1997; Tóth et al., 1997). Therefore, the GABAergic SH pathway has been proposed to be responsible for producing the correct levels of excitation, as well as regulating synchronous neuronal activities, including theta and gamma oscillations, which are crucial for memory and cognition (Colgin and Moser, 2010; Gangadharan et al., 2016; Hangya et al., 2009; Vertes, 2005).

Several studies have reported alterations in the GABAergic SH pathway associated with AD. For instance, J20 mice expressing human amyloid precursor protein (hAPP) with familial AD Swedish and Indiana mutations (Mucke et al., 2000) present a marked deterioration, displayed as a reduced number and complexity of GABAergic SH axon terminals; this decline also correlates with electrophysiological changes within the hippocampal network (Rubio et al., 2012; Vega-Flores et al., 2014). Similarly, VLW mice expressing human Tau (hTau) with three mutations related to frontotemporal dementia with parkinsonism linked to chromosome 17 (FTDP-17) (Lim et al., 2001) show a decrease in GABAergic SH innervation (Soler et al., 2017). Moreover, VLW mice present epilepsy and GABA_A_ receptor-mediated hyperexcitability (García-Cabrero et al., 2013).

Interestingly, several research groups have recently described the existence of individuals presenting the histopathological hallmarks of AD (namely Aβ plaques and P-Tau tangles) in the absence of cognitive impairment (Bjorklund et al., 2012; Briley et al., 2016; Erten-Lyons et al., 2009; Iacono et al., 2008; Singh et al., 2020; Zolochevska et al., 2018). This clinical entity, Non-Demented with AD Neuropathology (NDAN), is thought to have intrinsic mechanisms conferring neuroprotection against the classical degenerative AD processes. Several factors have been proposed as responsible for this protection, such as increased neurogenesis (Briley et al., 2016), increased hippocampal and total brain volume (which could indicate a larger cognitive reserve) (Erten-Lyons et al., 2009), neuronal and nuclear hypertrophy (Iacono et al., 2008), and preservation of the essential elements of the synaptic machinery that are dysfunctional in AD (Bjorklund et al., 2012; Singh et al., 2020). It has also been reported that NDAN individuals have a differential gene expression pattern at the postsynaptic density, which differs from that of AD and control subjects and could contribute to the mechanisms that confer synapse protection against AD pathology (Zolochevska et al., 2018). The lack of animal models mimicking NDAN hinders its characterization and the search for the mechanisms involved in preventing cognitive decline.

To assess the synergic or opposing effects of Aβ and P-Tau on hippocampal neuron physiology, hippocampal activity rhythms, and cognition, we crossed J20 (hAPP^Sw,Ind^) and VLW (hTau^VLW^, hTau with mutations G272V, P301L, and R406W) mice to generate a double transgenic mouse model with both Aβ and Tau pathologies, which are characteristic of AD. The Aβ plaque load in the resulting J20/VLW mice did not differ from that present in single transgenic J20 animals. The analysis of Tau phosphoepitopes revealed high levels of Tau phosphorylated at residues Thr231 (pThr231) and Thr205 (pThr205) in *Cornu Ammonis* (CA) 1 pyramidal neurons in J20/VLW mice, similar to single transgenic VLW animals. In contrast, J20/VLW mice presented higher densities of hippocampal interneurons accumulating pThr205 and pSer262 Tau than single transgenic VLW animals, thereby indicating an interneuron-specific modulation of Tau phosphorylation by Aβ. Surprisingly, GABAergic SH innervation on hippocampal interneurons in J20/VLW mice did not differ from that in control animals. Examination of hippocampal electrophysiology revealed partial recovery of hippocampal theta oscillations in J20/VLW animals, such oscillations being greatly affected in J20 and VLW mice. Furthermore, recognition memory deficits associated with Aβ accumulation were prevented in the double transgenic mouse model. These data render J20/VLW mice a suitable animal model to further understand the cognitive preservation in NDAN subjects and propose the maintenance of the GABAergic SH pathway as a mechanism underlying cognitive neuroprotection in NDAN.

## MATERIALS AND METHODS

### Animals

For the histological procedures, control wild-type (WT) adult male mice (C57BL/6J strain; 8-month-old (mo) and 12 mo; *n* = 4–5 per group) and transgenic adult male littermates from three different lines with the same genetic background were used. A double transgenic mouse line was generated by crossing J20 and VLW mice, and the resulting double mutant J20/VLW animals (8 mo and 12 mo; *n* = 4–5 per group), as well as the single mutant hemizygous J20 (8 mo and 12 mo; *n* = 4 per group) and heterozygous VLW mice (8 mo; *n* = 3–4 per group), were used. J20 animals overexpress hAPP^Sw,Ind^, hAPP carrying two familial AD mutations, namely Swedish (K670N/M671L) and Indiana (V717F), under the control of the platelet-derived growth factor subunit β promoter. VLW mice overexpress hTau^VLW^, hTau with four tubulin-binding repeats and three mutations related to FTDP-17 (G272V, P301L, and R406W) under the control of the Thy-1 promoter. For the electrophysiological study, 8 mo WT, VLW, and J20/VLW mice were used (*n* = 4 per group). For the behavioral tests, 8 mo WT (*n* = 9), J20 (*n* = 6), VLW (*n* = 7), and J20/VLW (*n* = 5) mice were used. For all experiments, *n* expresses the number of individual animals.

Animals were kept on a 12 h light/dark schedule with access to food and water *ad libitum*. Experimenters were blinded to the genotype of mice until data acquisition was completed. All experiments were performed in accordance with the European Community Council Directive and the National Institute of Health Guide for the Care and Use of Laboratory Animals and were approved by the local ethical committees.

### Detection of SH fibers

Animals were anesthetized with a 10/1 mixture of Ketolar® (50 mg/mL ketamine chlorhydrate, Parke-Davis)/Rompun® (2 % xylidine-thiazine chlorhydrate, Bayer) and stereotaxically injected with an anterograde tracer, 10 % biotinylated dextran-amine (BDA; 10,000 MW, Molecular Probes), in the MSDB complex. Each animal received midline injections of the tracer into the MSDB complex at one anteroposterior (AP) level, and at two dorsoventral (DV) injection sites by iontophoresis. Stereotaxic coordinates in millimeters were (from Bregma): AP +0.7 and DV − 3.0 and − 3.7 (Paxinos and Franklin, 2001). This protocol results in intense BDA labeling in the MSDB complex, which contains the highest proportion of GABAergic SH neurons (Pascual et al., 2004). After 5− 6 days, animals were anesthetized as before and perfused with 4 % paraformaldehyde in 0.1 M phosphate buffer. Brains were post-fixed for 48 h in 4 % paraformaldehyde, cryoprotected in phosphate-buffered saline with 30 % sucrose, frozen, and 30-μm coronal sections were cut and stored in a cryoprotectant solution (30 % glycerol, 30 % ethylene glycol, 40 % 0.1 M phosphate buffer) at − 20 °C until use.

### Immunodetection

Sections from iontophoretically injected animals were processed for double immunodetection of BDA and interneuron markers (Pascual et al., 2004). Sections were incubated simultaneously with the ABC complex (Vector Laboratories; 1/100), to visualize BDA, and rabbit polyclonal antibodies against PV (Swant®; 1/3000) or glutamic acid decarboxylase isoforms 65 and 67 (GAD, Chemicon International; 1/2000), to visualize interneuron populations. BDA was developed with H_2_O_2_ and diaminobenzidine (DAB), nickel ammonium sulfate (Ni), and cobalt chloride (Co), yielding a black end product in SH fibers. Primary antibodies were visualized by sequential incubation with biotinylated secondary antibodies and the ABC complex (Vector Laboratories). Peroxidase activity was developed with H_2_O_2_ and DAB to produce a brown end product. Sections were mounted onto gelatinized slides, dehydrated, and coverslipped with Eukitt® (O. Kindler).

To detect pThr231, pThr205, or pSer262 Tau, sections were incubated with AT-180 mouse anti-phosphothreonine 231 (Innogenetics; 1/300), T205 rabbit anti-phosphothreonine 205 (Invitrogen™; 1/1000), or S262 rabbit anti-phosphoserine 262 (Invitrogen™; 1/100) antibodies. Subsequent steps were performed as described.

For the detection of Aβ plaques, sections were incubated with 3D6 mouse anti-Aβ (amino acids 1–5) antibody (obtained from the supernatant of cultured Murine Hybridoma Cell Line, RB96 3D6.32.2.4 (PTA-5130), American Type Culture Collection; 1/200). Subsequent steps were performed as described.

To detect GABAergic SH projection neurons, sections corresponding to the medial septum were incubated with rabbit anti-PV antibody. Primary antibody was visualized by sequential incubation with biotinylated secondary antibody and the ABC complex (Vector Laboratories). Subsequent steps were performed as described.

To determine whether hippocampal interneurons accumulate P-Tau, double fluorescent immunodetections were conducted using AT-180, T205, or S262 and PV, Calretinin (CR), or Calbindin (CB) primary antibodies. Sections were incubated simultaneously with goat anti-PV (Swant®; 1/3000), goat anti-CR (Swant®; 1/3000), or mouse anti-CB (Swant®; 1/3000) antibodies and AT-180, T205, or S262. They were then incubated with Alexa Fluor 568 donkey anti-goat or anti-mouse IgG against interneuron primary antibodies, whereas P-Tau primary antibodies were targeted with Alexa Fluor 488 donkey anti-rabbit IgG or anti-mouse IgG (Invitrogen™; 1/1000). Sections were mounted onto slides and coverslipped with Mowiol® (Merck).

### Analysis of histological sections

Microscopic observations focused on sections corresponding to the medial septum and to dorsal (sections between AP − 1.6 and − 2.3 mm from Bregma) and ventral (sections between AP − 2.9 and − 3.4 mm from Bregma) hippocampal levels, following the atlas reported by Paxinos and Franklin (Paxinos and Franklin, 2001).

To estimate the density of hippocampal interneurons and the percentage of these cells contacted by GABAergic SH fibers, the density of interneurons containing GAD or PV and the percentage of these receiving BDA-positive pericellular baskets was calculated in distinct regions of the hippocampal area (CA1, CA3, and dentate gyrus (DG)) of each section (8 mo and 12 mo WT and J20/VLW mice; *n* = 4–5 animals per group, 3 sections per animal). The area comprising the hippocampal region of each section was quantified using Fiji software (Schindelin et al., 2012). Due to the large number of GAD-immunopositive cells, several sample areas were selected for each section (8 mo and 12 mo WT and J20/VLW mice; *n* = 4 animals per group, 3 sections per animal). The selected samples (125 µm-wide stripes) contained all hippocampal layers (perpendicularly from the ventricle to the pial surface), and each section included the CA1, the CA3, and the DG. Data were represented as density of interneurons per square millimeter and percentage of GAD- or PV-positive cells contacted by GABAergic SH fibers. To assess the complexity of GABAergic SH contacts, synaptic boutons around the soma of GAD- or PV-positive cells were counted in the same sample areas under an optical microscope (Nikon E600, Nikon Corporation), and data were expressed as number of boutons per basket.

To estimate the density of interneurons accumulating pThr231, pThr205, or pSer262 Tau, and to compute the mean gray value of the pThr231 and pThr205 signals in pyramidal neurons, samples immunodetected with AT-180, T205, or S262 antibodies were scanned with a NanoZoomer 2.0HT whole slide imager (Hamamatsu Photonics) at 20x. The density of pThr231, pThr205, and pSer262 Tau-immunopositive interneurons was quantified in distinct regions of the hippocampal area (CA1, CA3, and DG) of each section (8 mo WT, J20, VLW, and J20/VLW mice; *n* = 4 animals per group, 3 sections per animal) and data were expressed as density of cells per square millimeter. The cells and area comprising each hippocampal region were quantified using Fiji software (Schindelin et al., 2012). The mean gray value of the pThr231 and pThr205 Tau signals was calculated in 10 square millimeter stripes in matching regions of the pyramidal layer of the CA1 after thresholding the images to exclude background signal (8 mo VLW and J20/VLW mice; *n* = 3–4 animals per group, 3 sections per animal) using Fiji software (Schindelin et al., 2012).

To assess the percentage of Aβ plaques in the hippocampus, samples immunodetected with 3D6 antibody were scanned with a NanoZoomer 2.0HT whole slide imager (Hamamatsu Photonics) at 20x. The Trainable Weka Segmentation plugin (Arganda-Carreras et al., 2017) from Fiji software (Schindelin et al., 2012) was applied to the images by using a set of machine-learning algorithms with a collection of image features selected by the user to produce pixel-based segmentations. Images were processed using a macro provided by Sebastién Tosi (Institute for Research in Biomedicine, Barcelona) to identify and quantify Aβ plaques (8 mo and 12 mo J20 and J20/VLW mice; *n* = 4 animals per group, 3 sections per animal).

To estimate the density of GABAergic SH neurons, samples stained with PV were scanned with a NanoZoomer 2.0HT whole slide imager (Hamamatsu Photonics) at 20x. The density of GABAergic SH neurons was quantified in serial MSDB complex sections (8 mo WT and J20/VLW mice; *n* = 4 animals per group, 4 sections per animal) and was defined as the density of PV-immunopositive cells per square millimeter. The cells and the area comprising the MSDB complex were quantified using Fiji software (Schindelin et al., 2012).

### Statistical analysis of histological data

Histological data were processed for statistical analysis with GraphPad Prism 8 (GraphPad Software Incorporated). All data were tested for normal distribution. To examine differences between two experimental groups, unpaired two-tailed Student’s *t*-test or Welch’s *t*-test were used, when the samples had equal variances or not, respectively. To assess differences between more than two experimental groups, one-way ANOVA was used. Post hoc comparisons were performed by Tukey’s test only when a significant main effect of one-way ANOVA was revealed. Significance level was set at *p* < .05: \**p* < .05, \*\**p* < .01. Statistical values are presented as mean ± standard error of the mean (SEM).

### Image acquisition

Optical microscopy (Nikon E600, Nikon Corporation) images of the immunohistochemically-stained brain sections were acquired through a digital camera (Olympus DP72, Olympus Corporation) coupled to the microscope and were processed by Cell F^ software (Olympus Corporation).

Confocal microscopy (Leica TCS SP5, Leica Microsystems) images of the immunofluorescence-stained samples were acquired using LAS AF software (Leica Microsystems). To observe possible colocalization between the interneuron and P-Tau markers, images were processed by Fiji software (Schindelin et al., 2012).

### Surgery for the chronic recording of hippocampal local field potentials

Eight mo WT, VLW, and J20/VLW mice were anesthetized with 0.8–3 % halothane delivered from a calibrated Fluotec 5 vaporizer (Datex-Ohmeda) at a flow rate of 1–2 L/min oxygen. Animals were implanted with a recording electrode aimed at the ipsilateral *stratum radiatum* underneath the hippocampal CA1 area. Stereotaxic coordinates in millimeters were (from Bregma): AP − 2.2, lateromedial (LM) 1.2, and DV − 1.0 to − 1.5 (Paxinos and Franklin, 2001). These electrodes were made of 50-µm Teflon-coated tungsten wire (Advent Research Materials Ltd). A bare silver wire (0.1 mm) was affixed to the skull as a ground. All wires were soldered to a 6-pin socket, and the socket was fixed to the skull with the help of two small screws and dental cement (see (Gruart et al., 2006) for details).

### Recording procedures

The electroencephalographic field activity of the CA1 area was recorded with the help of Grass P511 differential amplifiers. LFP recordings were carried out with the awake animal placed in either a small box (5 x 5 x 5 cm) to prevent walking movements or a large box (20 x 15 x 15 cm) in which the animal could move freely. Recordings were carried out for 20 min, of which up to 5 min of recording free of unwanted artifacts were selected for spectral analysis.

### Data analysis of electrophysiological studies

Hippocampal activity was stored digitally on a computer through an analog/digital converter (CED 1401 Plus, CED), at a sampling frequency of 11–22 kHz and an amplitude resolution of 12 bits. Commercial computer programs (Spike 2 and SIGAVG, CED) were modified to represent recorded LFPs.

The power spectrum of hippocampal LFPs collected during recording sessions was computed with the help of Mat Lab 7.4.0 software (MathWorks), using the fast Fourier transform with a Hanning window, expressed as relative power and averaged across each session. This average was analyzed and compared using the wide-band model, considering the following bands: delta (< 4 Hz), theta (4.1–8 Hz), alpha (8.1–12 Hz), beta (12.1–26 Hz), and gamma (26.1–100 Hz) (Fernández-Lamo et al., 2016; Múnera et al., 2000).

### Behavioral tests

#### Novel object-recognition test (NORT)

Object-recognition memory was assessed following a protocol previously described (Puighermanal et al., 2009). Briefly, on day one, mice were habituated to a V-shaped maze (each corridor measuring 30 cm long × 4.5 cm wide × 15 cm high) for 9 min. On day two, two identical objects (familiar objects) were located at the end of each corridor for 9 min, and the time that the mice spent exploring each object was measured. Twenty-four hours later, one of the familiar objects was replaced by a new object (novel object) (see Fig. 1A). The time spent exploring each of the objects was computed to calculate a discrimination index (DI). The DI was calculated as the difference between the time spent exploring the novel object (Tn) minus the time exploring the familiar object (Tf) divided by the total exploration time (addition of the time exploring both objects), (DI = (Tn – Tf) / (Tn + Tf)). Object exploration was defined as the orientation of the nose towards the object at a distance of < 2 cm. Mice that explored both objects for < 10 s or one object for < 3 s were excluded from the analysis. A higher DI is considered to reflect greater memory retention for the familiar object. Total exploration time was considered a measure of general activity during the test.

**Figure 1.**
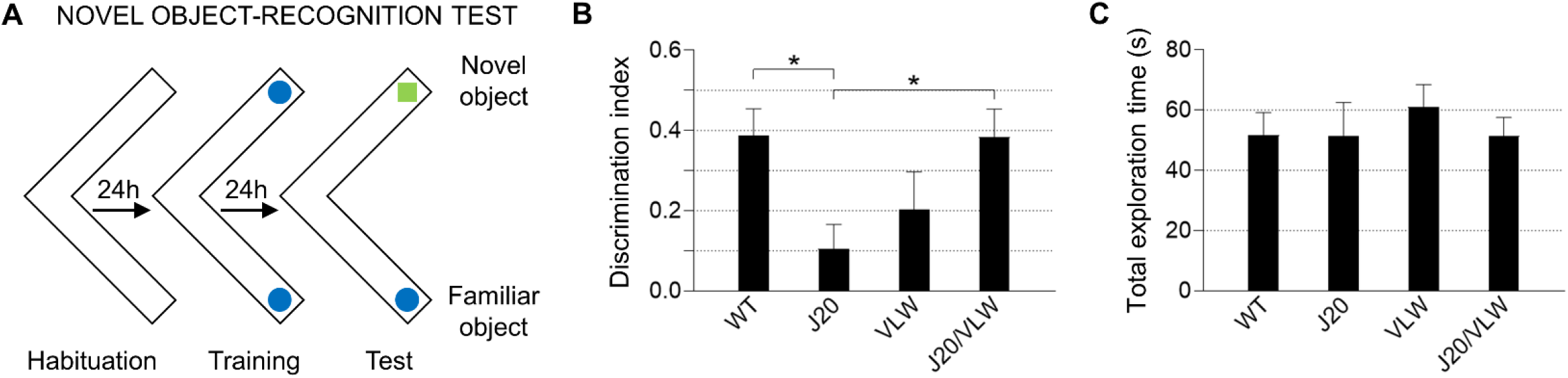
No recognition memory deficits are present in J20/VLW mice. **(A)** Protocol for novel object-recognition test. The time spent exploring each of the objects during the test phase was computed to calculate a discrimination index. **(B)** Discrimination index in novel object-recognition test. A higher discrimination index is considered to reflect greater memory retention for the familiar object. **(C)** Total exploration time in novel object-recognition test was considered a measure of general activity. For (B) and (C): one-way ANOVA, \**p* < .05. All behavioral tests were performed on 8 mo WT (*n* = 9), J20 (*n* = 6), VLW (*n* = 7), and J20/VLW (*n* = 5) mice. Error bars represent SEM.

#### Elevated plus-maze

Anxiety-like behavior was evaluated using the elevated plus-maze test. The setup consisted of a black Plexiglas apparatus with four arms (29 cm long x 5 cm wide)—two open and two closed—set in a cross from a neutral central square (5 x 5 cm) elevated 40 cm above the floor. Light intensity in the open and closed arms was 45 and 5 luxes, respectively. Mice were placed in the central square facing one of the open arms and tested for 5 min. The percentage of entries into the open arms was determined as 100 x (entries into open arms) / (entries into open arms + entries into closed arms). Animals that exited the maze during exploration were excluded from the analysis. Total entries into each arm were calculated as a control of exploratory behavior.

#### Rotarod

Motor coordination was assessed using the accelerating rotarod (5-lane accelerating rotarod; LE 8200, Panlab). On day one, mice were trained to hold onto the rod at a constant speed (4 rpm) for at least 120 s. On day two, mice were trained to hold onto the rod at a constant speed higher than the previous day (6 rpm) for at least 120 s. On day three, the test was performed. During the test, the rod accelerated from 4 to 40 rpm within 1 min, and latency to fall and maximum speed were measured in five consecutive trials. Data are expressed as the mean of the five trials.

### Statistical analysis of behavioral tests

Behavioral data were processed for statistical analysis with GraphPad Prism 8 (GraphPad Software Incorporated). Comparisons between experimental groups were performed by one-way ANOVA. Post hoc comparisons were performed by Fisher’s LSD test only when a significant main effect of one-way ANOVA was revealed. Significance level was set at *p* < .05. Statistical values are presented as mean ± SEM.

## RESULTS

### Cognitive recovery of J20/VLW mice in contrast to J20 animals

J20 and VLW mice show considerable cognitive deficits, mainly in spatial memory (Cissé et al., 2011; Harris et al., 2010; Navarro et al., 2008). To characterize double transgenic J20/VLW mice, which accumulate both Aβ and P-Tau, we performed various tests to analyze long-term memory, comparing 8 mo WT, J20, VLW, and J20/VLW animals. First, we determined that there were no major differences in either locomotion or anxiety between the four experimental groups (Supplementary Fig. 1). Surprisingly, alterations in NORT performance of J20 animals, which have been previously described (Cissé et al., 2011; Harris et al., 2010) were not present in J20/VLW mice, which showed a behavior equivalent to that of WT mice (Fig. 1B). These data suggest that the specific forms of P-Tau in VLW mice, in coexistence with Aβ accumulation, protect against the recognition memory deficits associated with Aβ. Such specific differences were not related to exploratory behavior or motility, since total exploration times in the NORT were similar in the four experimental groups (Fig. 1C).

### Hippocampal rhythmic activity preservation in J20/VLW mice

Next, we analyzed the oscillatory activities of hippocampal circuits in 8 mo behaving WT, J20, VLW, and J20/VLW mice. Chronically implanted electrodes recorded hippocampal field activity in behaving animals placed in either small or large boxes, to determine the contribution of overt motor activities to the power spectra of the theta and gamma bands (Fig. 2A, B). As previously described (Rubio et al., 2012), the spectral analysis of LFP recordings showed a clear decrease in the spectral power of the theta band in J20 mice placed in small and large boxes (43 % and 38 %, respectively) compared to age-matched WT animals (Fig. 2C). In addition, a considerable reduction in theta spectral power was present in VLW animals placed in small and large boxes (33 % and 42 %, respectively). In contrast, J20/VLW mice showed only a slight decrease in theta spectral power when located in small and large boxes (8 % and 22 %, respectively). We also examined the spectral power of the gamma band (Fig. 2D). In a previous study (Rubio et al., 2012), we found a 50 % reduction in gamma spectral power in J20 animals placed in large and small boxes compared to WT mice. The results presented herein demonstrate that the gamma band in VLW animals was equivalent to that of WT mice. Interestingly, our data indicate a minor reduction (33 %) in this band in J20/VLW mice compared to age-matched WT animals. We conclude that both theta and gamma oscillations are rescued in J20/VLW mice. Thus, coexistence of Aβ and P-Tau preserve hippocampal rhythmic activity.

**Figure 2.**
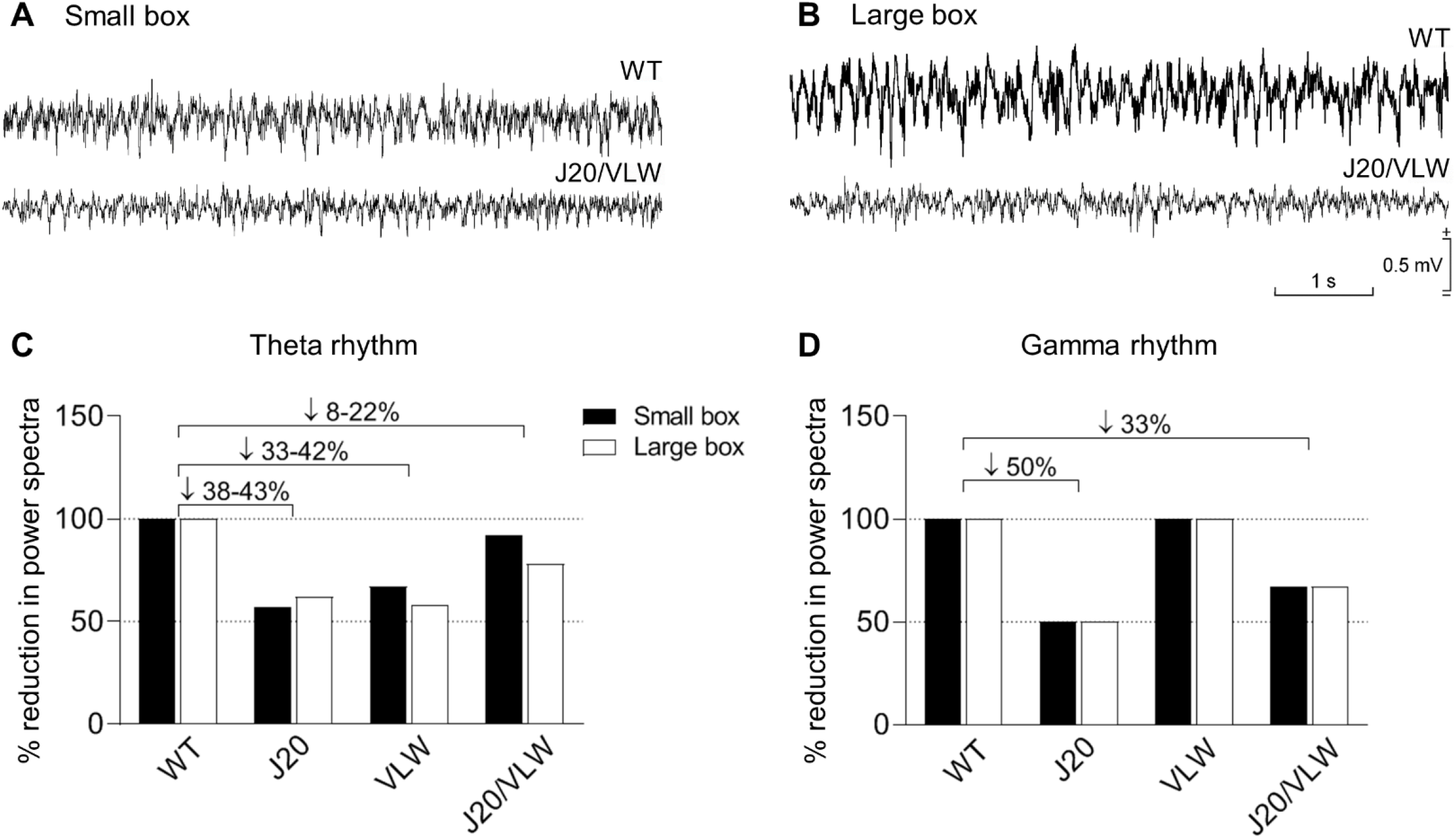
Hippocampal oscillations, which are markedly impaired in J20 and VLW animals, are partially rescued in J20/VLW mice. **(A and B)** Representative examples of LFP activity recorded in the CA1 region of the hippocampus of 8 mo WT and J20/VLW mice located in either a small (A) or a large (B) box. **(C and D)** Spectral powers were computed from similar records collected from 8 mo WT, J20, VLW, and J20/VLW mice. Histograms representing the percent decrease in spectral power for theta (C) and gamma (D) bands, comparing J20, VLW, and J20/VLW mice to WT animals. *n* = 4 animals per group.

### No changes in Aβ accumulation in J20/VLW animals compared to J20 mice

To study whether the presence of P-Tau induces changes in Aβ accumulation, we performed immunodetection with 3D6 antibody (which specifically detects amino acids 1–5 of Aβ) on J20/VLW and J20 hippocampal sections. Quantification of the percentage of Aβ plaques confirmed no changes in the accumulation of Aβ plaques in the hippocampus of J20/VLW mice compared to J20 animals at 8 mo (Fig. 3A-C). Given that the amount of plaques is minor at this maturation stage, we then analyzed 12 mo animals. The percentage of Aβ plaques in the hippocampus of 12 mo J20/VLW and J20 littermates was similar (Fig. 3D-F), suggesting that the cognitive and physiological improvements observed in J20/VLW animals is not a consequence of a reduction in Aβ plaque load in double transgenic mice.

**Figure 3.**
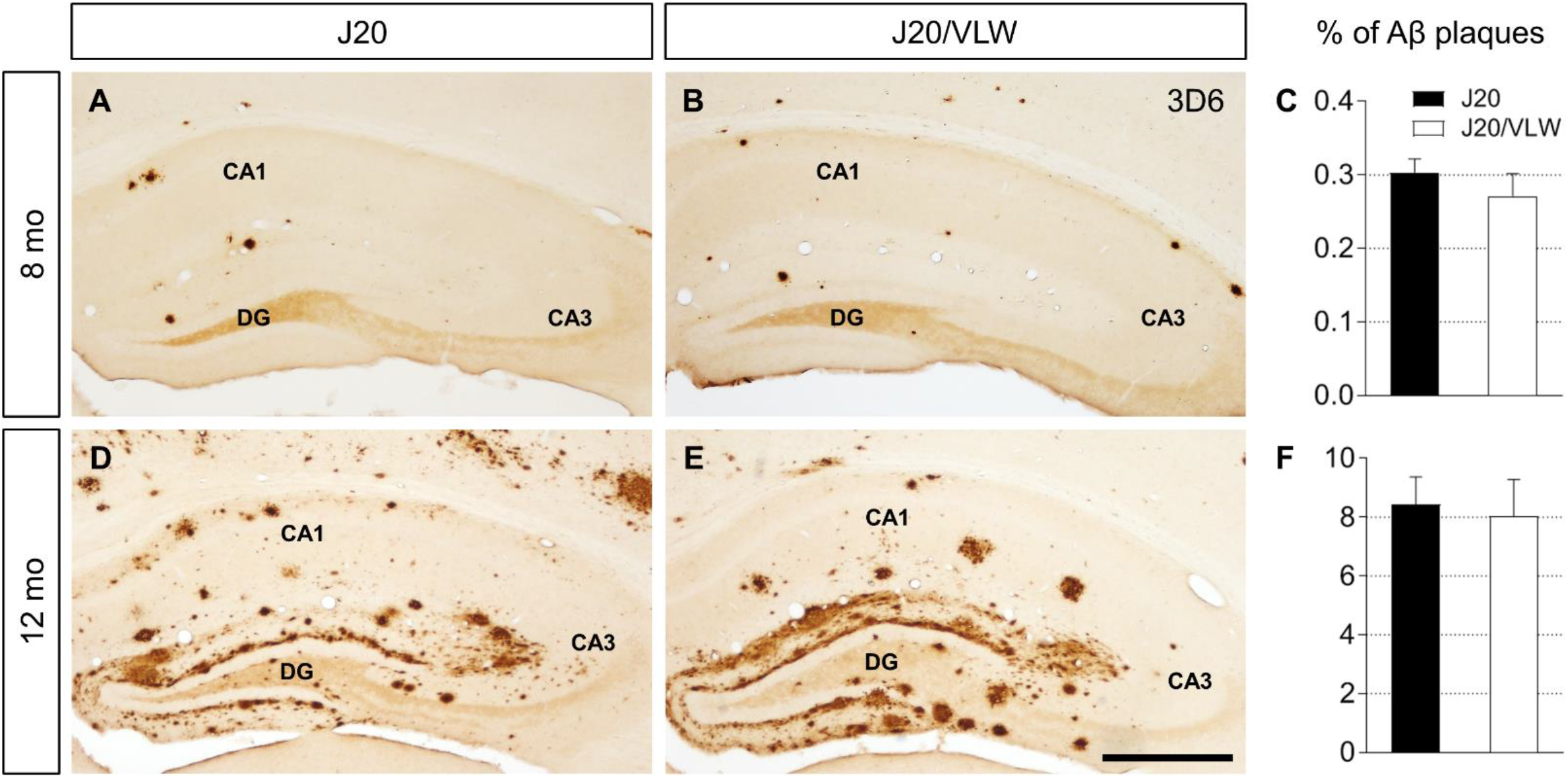
Aβ plaque load of J20/VLW mice is similar to that of J20 animals. Immunodetection of Aβ plaques with 3D6 antibody in hippocampal sections from 8 mo and 12 mo J20 and J20/VLW mice. **(A, B, D, and E)** Accumulation of Aβ plaques in the hippocampus of 8 mo (A and B) and 12 mo (D and E) J20 and J20/VLW mice. **(C and F)** Quantification of the percentage of Aβ plaques in the hippocampus of 8 mo (C) and 12 mo (F) J20 and J20/VLW animals. For (C) and (F): Student’s *t*-test. *n* = 4 animals per group, 3 sections per animal. Error bars represent SEM. Scale bar: 500 µm.

### Differential Tau phosphorylation signature in hippocampal GABAergic interneurons of J20/VLW mice

Aβ has been described to induce Tau phosphorylation both *in vivo* and *in vitro* (Jin et al., 2011; Zempel et al., 2010). Thus, we examined the pattern of Tau phosphorylation by immunodetection. We show that there are no changes in the accumulation of Tau phosphorylated at residues Thr231 or Thr205 in the somatodendritic compartment of pyramidal neurons when comparing VLW and J20/VLW mice (Fig. 4A-D). No differences in the pThr231 and pThr205 Tau signals in the CA1 pyramidal layer were detected between 8 mo VLW and J20/VLW animals (Fig. 4C, D). No pSer262 Tau was detected in the pyramidal layer of either VLW or J20/VLW mice.

**Figure 4.**
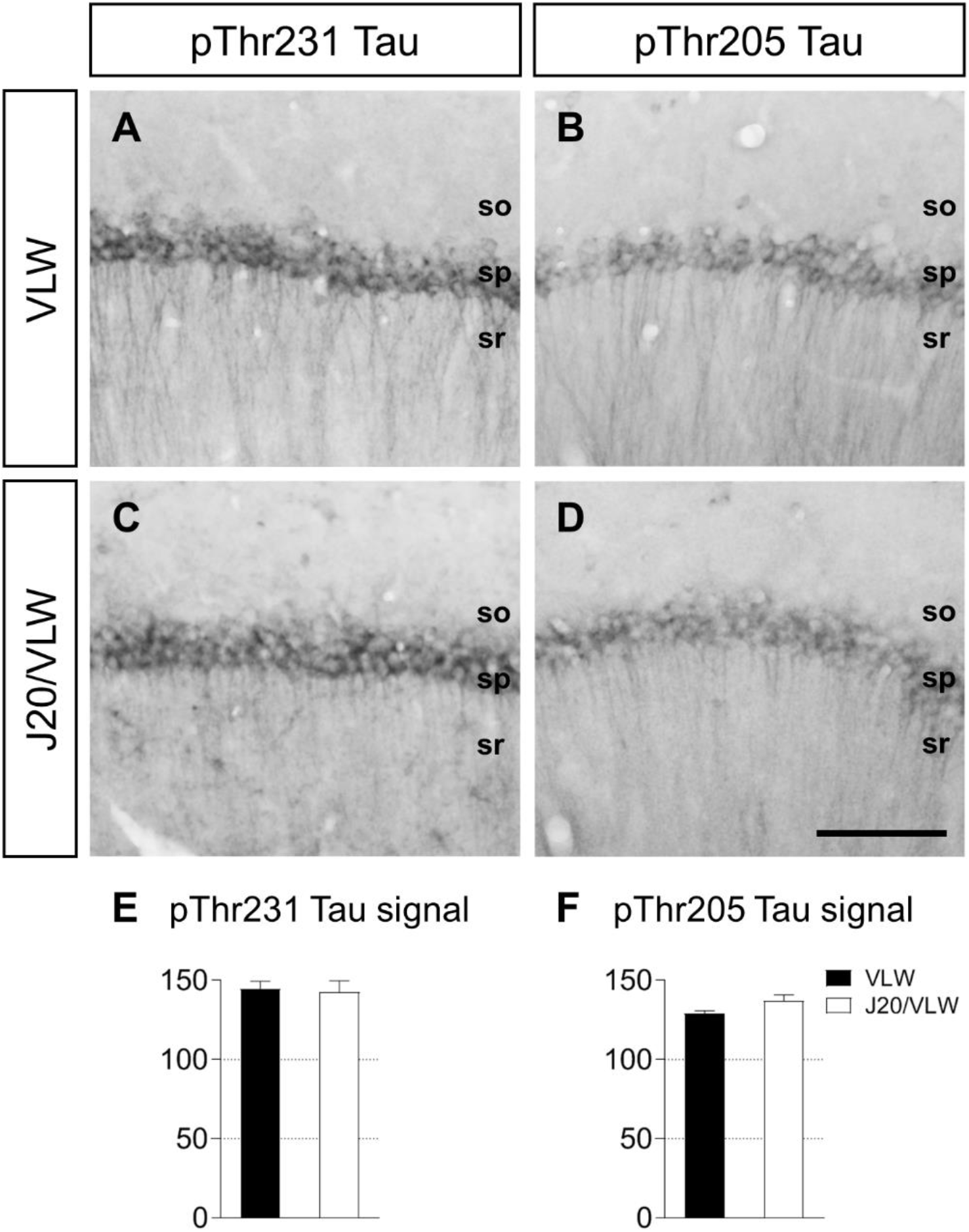
Pyramidal neurons of VLW and J20/VLW mice accumulate similar amounts of pThr231 and pThr205 Tau. Immunodetection and quantification of pThr231 and pThr205 Tau-positive signal in hippocampal sections from 8 mo VLW and J20/VLW mice. **(A-D)** pThr231 (A and C) and pThr205 Tau (B and D) accumulation in pyramidal neurons in the CA1 region of the hippocampus of VLW and J20/VLW mice. **(E and F)** Quantification of the mean gray value of pThr231 (E) and pThr205 Tau (F) signal in the pyramidal layer of VLW and J20/VLW animals. For (E) and (F): Student’s *t*-test. *n* = 3–4 animals per group, 3 sections per animal. Error bars represent SEM. Abbreviations: so, *stratum oriens*; sp, *stratum pyramidale*; sr, *stratum radiatum*. Scale bar: 100 µm.

We next compared the distribution and density of interneurons accumulating P-Tau in VLW and J20/VLW mice, as recent data reports the presence of pThr231, pThr205, and pSer262 Tau in the somas of hippocampal interneurons in VLW mice (Dávila-Bouziguet et al., 2019). Our data indicate that some hippocampal interneurons located mainly in the *stratum oriens* of the CA1 accumulate pThr231 Tau both in VLW and J20/VLW mice (Fig. 5A, D). Although no statistical differences in the density of pThr231 Tau-positive interneurons were found between the VLW and J20/VLW hippocampus, a clear upward trend was observed in double transgenic mice (Fig. 5G). Subsequently, we examined the density and distribution of pThr205 Tau-positive cells. In agreement with our previous studies, we found that Aβ in J20 mice induced an increase in the density of pThr205 Tau-positive hippocampal interneurons, in comparison with WT mice, and that P-Tau accumulation in VLW mice induced a reduction in the pThr205 Tau-positive interneuron density (Fig. 5B, H) (Dávila-Bouziguet et al., 2019). In J20/VLW double transgenic mice, the density of pThr205 Tau-positive GABAergic interneurons was higher than in VLW animals and lower than in J20 mice, being similar to that of WT animals (Fig. 5E, H). Thus, our data indicate that the increase in the density of pThr205 Tau-positive interneurons induced by Aβ and the decrease observed in the VLW hippocampus are abolished by the simultaneous presence of Aβ and P-Tau in J20/VLW animals (Fig. 5B, E, and H). Finally, we show a considerable increase in the density of interneurons accumulating pSer262 Tau in J20/VLW mice compared to VLW animals (Fig. 5C, F, and I). This observation suggests a synergic effect of P-Tau and Aβ in the induction of Tau phosphorylation at residue Ser262 in hippocampal interneurons. We thus conclude that coexistence of Aβ and P-Tau in J20/VLW mice confers a particular Tau phosphorylation signature in GABAergic hippocampal interneurons, but not so in hippocampal principal neurons.

**Figure 5.**
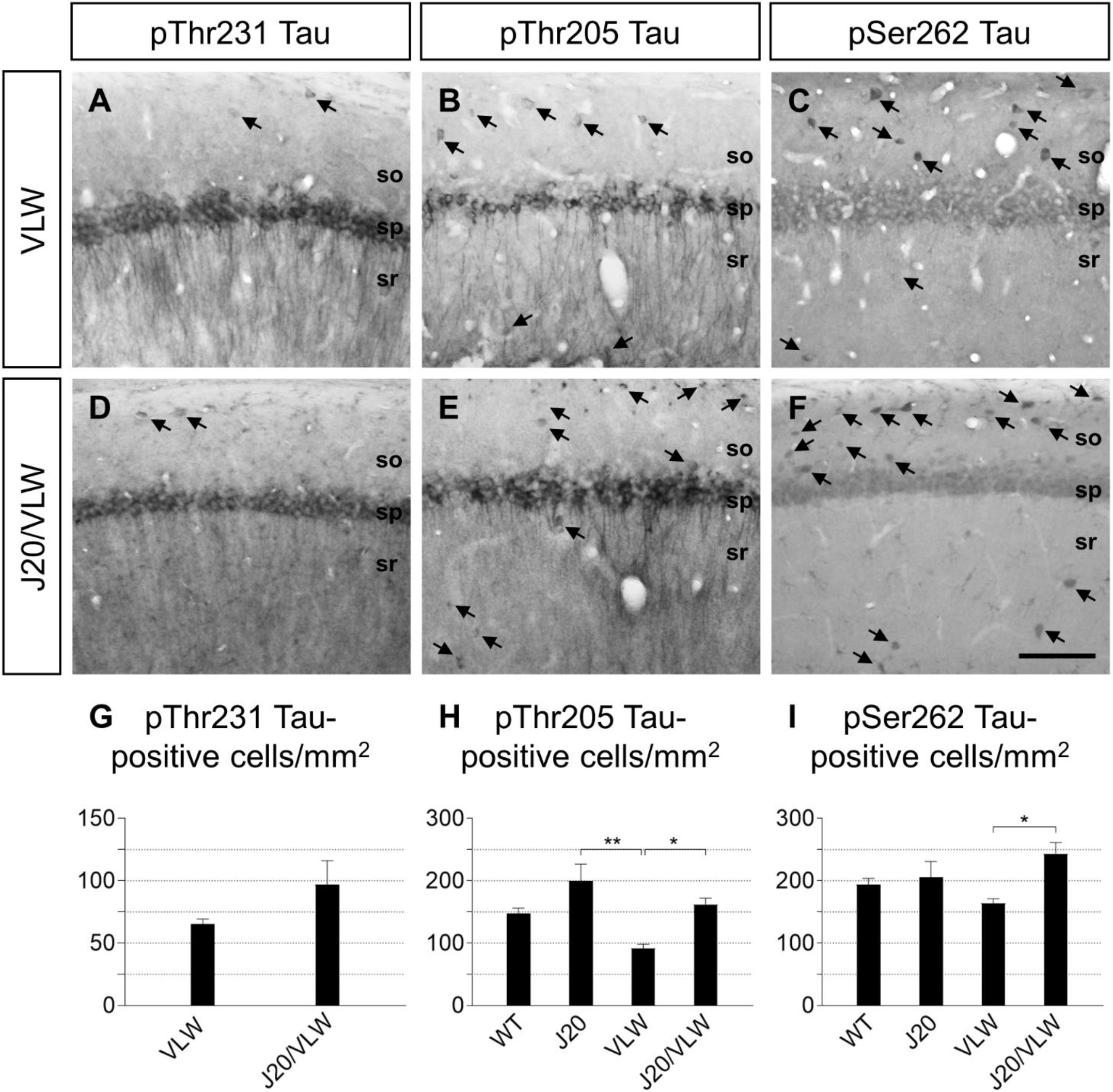
Increased density of hippocampal interneurons accumulating P-Tau in J20/VLW mice, which accumulate Aβ peptide, compared to VLW animals. Immunodetection and cell density quantification of pThr231, pThr205, and pSer262 Tau-positive cells in hippocampal sections from 8 mo WT, J20, VLW, and J20/VLW mice. **(A-F)** Accumulation of pThr231 (A and D), pThr205 (B and E), and pSer262 Tau (C and F) in hippocampal interneurons (arrows) in the CA1 region of the hippocampus of VLW and J20/VLW mice. **(G)** Density quantification of pThr231 Tau-positive interneurons in the hippocampus of VLW and J20/VLW mice. **(H and I)** Density quantification of pThr205 (H) and pSer262 (I) Tau-positive interneurons in the hippocampus of WT, J20, VLW, and J20/VLW mice. For (G): Welch’s *t*-test. For (H) and (I): one-way ANOVA, \**p* < .05, \*\**p* < .01. *n* = 4 animals per group, 3 sections per animal. Error bars represent SEM. Abbreviations: so, *stratum oriens*; sp, *stratum pyramidale*; sr, *stratum radiatum*. Scale bar: 100 µm.

To characterize the interneuron subtypes accumulating P-Tau, double fluorescent immunodetection of pThr231, pThr205, and pSer262 Tau, combined with detection of the interneuron markers PV, CR, and CB, was performed on J20/VLW hippocampal slices. As described in VLW animals, pThr231 Tau accumulated specifically in some PV-positive interneurons throughout the hippocampus (Supplementary Fig. 2A-C), while some PV-, CR-, and CB-positive interneurons scattered in different hippocampal layers and regions accumulated pThr205 and pSer262 Tau (Supplementary Fig. 2D-L). In agreement with previous data in human AD (Blazquez-Llorca et al., 2010), only a few PV-positive cells with pThr205 Tau in their soma were present in the J20/VLW hippocampus.

### GABAergic SH pathway preservation in J20/VLW mice

The GABAergic SH pathway is crucial for the activity of hippocampal interneurons and for hippocampal electrophysiology and cognition (Etter et al., 2019; Hangya et al., 2009). Given that the above data demonstrate the preservation of cognition and hippocampal rhythmic activities in double transgenic J20/VLW mice and alteration of the pattern of P-Tau specifically in interneurons, we next investigated whether the GABAergic SH network was preserved in these double transgenic mice. To map the GABAergic SH pathway, we performed stereotaxic injections of an anterograde tracer in the MSDB complex (Pascual et al., 2004; Rubio et al., 2012; Soler et al., 2017). In agreement with our previous studies, single transgenic J20 and VLW mice showed a 40 % and a 36 % reduction in the percentage of GABAergic hippocampal interneurons (GAD-immunopositive) innervated by GABAergic SH fibers, respectively, compared to WT animals (Fig. 6G) (Rubio et al., 2012; Soler et al., 2017). In addition, J20 animals showed a 34 % decrease in the number of GABAergic SH boutons on GAD-positive hippocampal cells (Fig. 6H) (Rubio et al., 2012). These findings suggest that the impairment of GABAergic SH innervation, which modulates hippocampal network activities, may be linked to the cognitive deficits present in J20 and VLW mice (Rubio et al., 2012; Soler et al., 2017). We next characterized the GABAergic SH pathway in J20/VLW animals. Neither the distribution nor percentage of GAD-immunopositive neurons contacted by septal GABAergic fibers nor the complexity (number of boutons per cell) of the GABAergic SH synaptic contacts were altered in 8 mo J20/VLW mice compared to WT animals (Fig. 6). We next studied the pattern GABAergic SH innervation on PV-positive hippocampal interneurons in J20/VLW animals, the hippocampal interneuron population most affected by the loss of GABAergic SH innervation in single transgenic J20 and VLW mice (Rubio et al., 2012; Soler et al., 2017). These single transgenic mice exhibited a 30 % and a 36 % reduction, respectively, in the percentage of PV-positive cells contacted by GABAergic SH fibers. Moreover, the complexity of the GABAergic SH contacts on PV-positive cells was reduced by 36 % and 34 % in J20 and VLW mice, respectively (Fig. 7G, H) (Rubio et al., 2012; Soler et al., 2017). In contrast, the distribution and the percentage of PV-containing interneurons contacted by GABAergic SH fibers were spared in J20/VLW mice, and only a slight decrease in the complexity of the GABAergic SH contacts on PV-positive cells occurs in double transgenic mice (Fig. 7).

**Figure 6.**
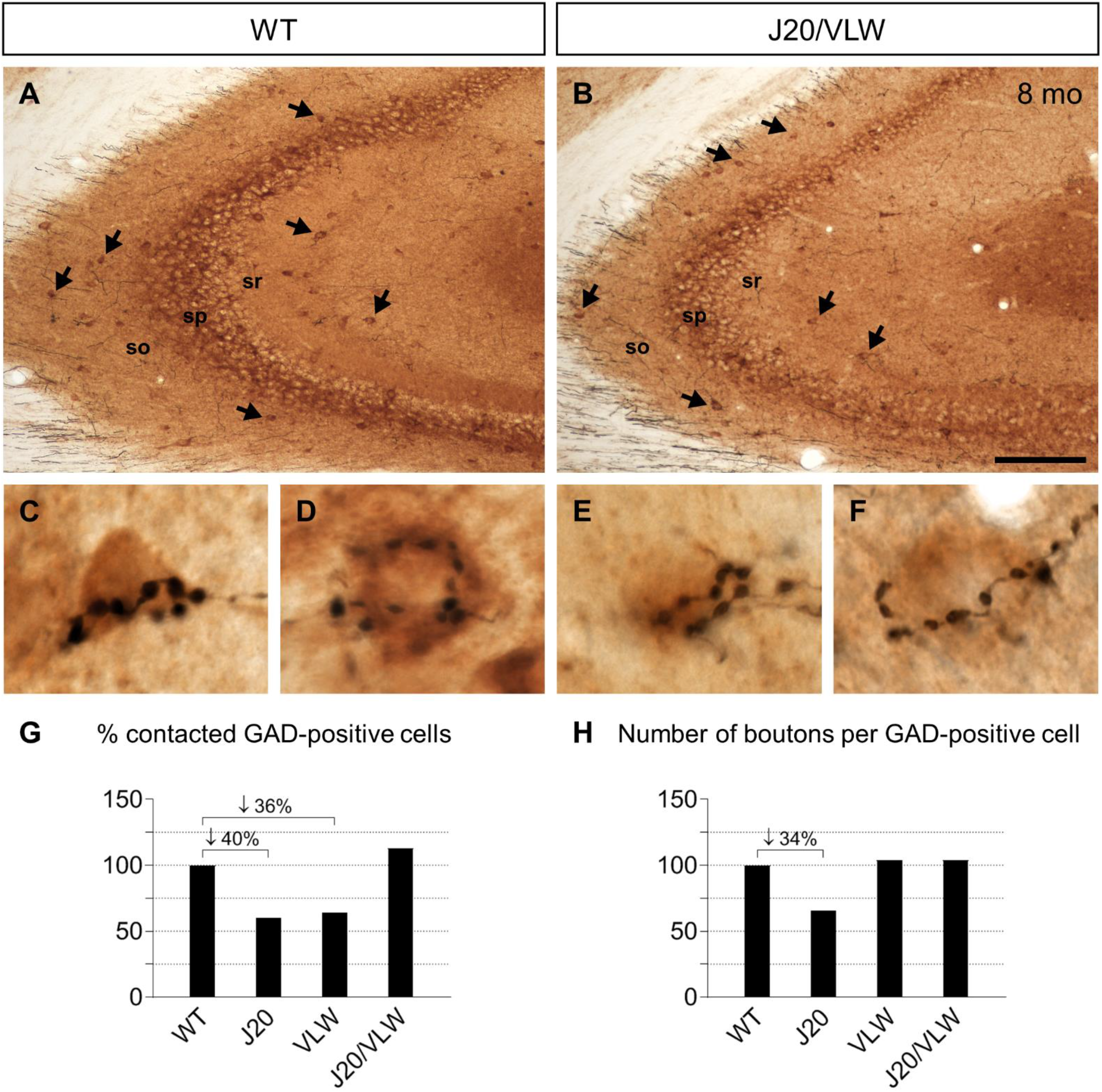
GABAergic SH innervation on GAD-positive cells is preserved in 8 mo J20/VLW mice. Double immunodetection of GABAergic SH fibers and GAD-positive cells in hippocampal sections from 8 mo WT and J20/VLW mice. **(A and B)** GABAergic SH fibers contacting GAD-positive cells (arrows) in the CA3 region of WT (A) and J20/VLW (B) mice. (C-F) GABAergic SH baskets forming synaptic boutons (black) on the soma of GAD-positive cells (brown) in WT (C and D) and J20/VLW (E and F) mice. **(G and H)** Representation of the percentage of change in the percentage of GAD-positive cells contacted by GABAergic SH fibers, and in the complexity of GABAergic SH contacts, comparing the four experimental groups. *n* = 4 animals per group, 3 sections per animal. Abbreviations: GAD, glutamic acid decarboxylase; so, *stratum oriens*; sp, *stratum pyramidale*; sr, *stratum radiatum*. Scale bar: 150 µm (A and B), 10 µm (C-F).

**Figure 7.**
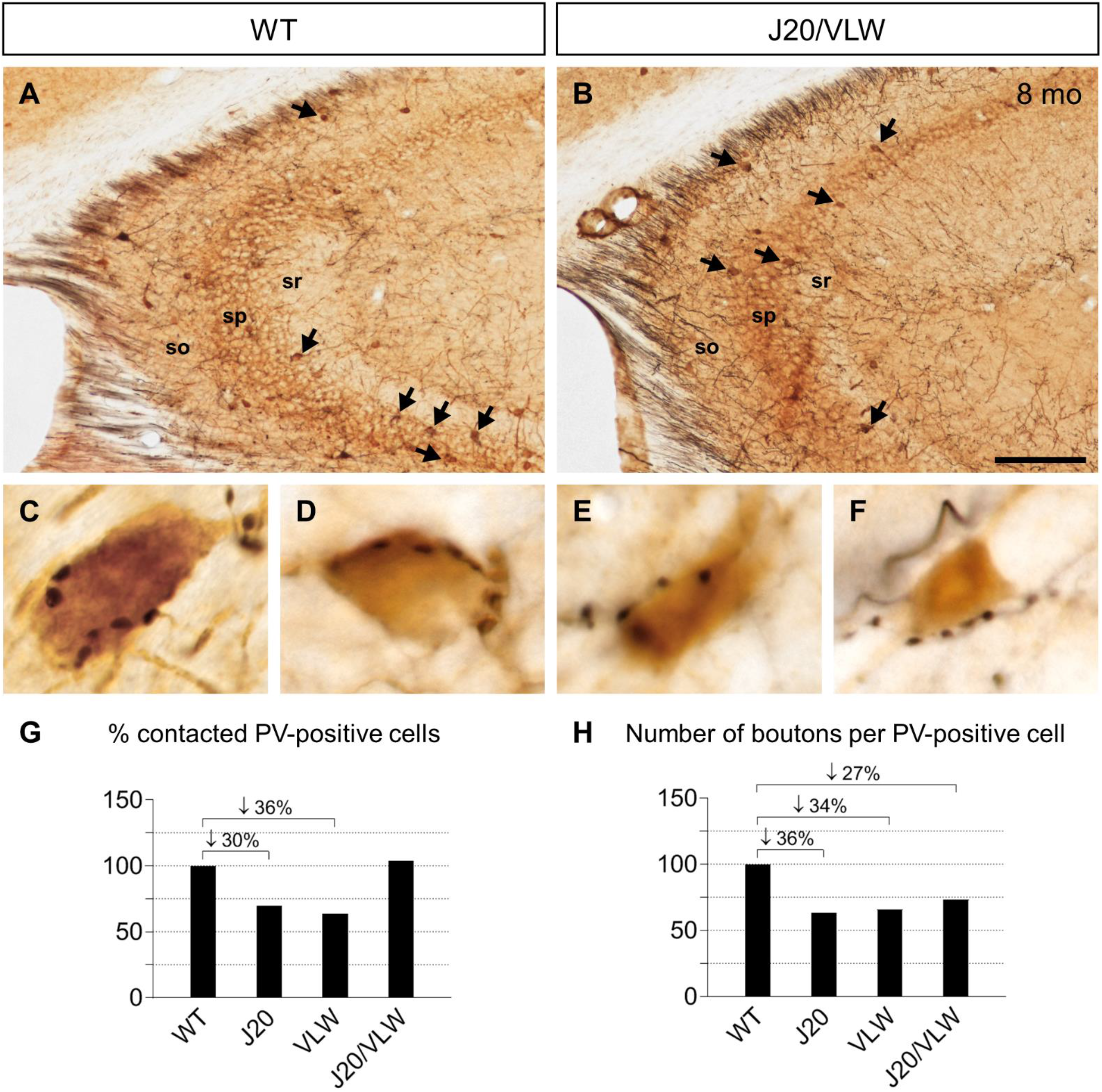
GABAergic SH innervation on PV-positive cells is preserved in 8 mo J20/VLW mice. Double immunodetection of GABAergic SH fibers and PV-positive cells in hippocampal sections from 8 mo WT and J20/VLW mice. **(A and B)** GABAergic SH fibers contacting PV-positive cells (arrows) in the CA3 region of WT (A) and J20/VLW (B) mice. **(C-F)** GABAergic SH baskets forming synaptic boutons (black) on the soma of PV-positive cells (brown) in WT (C and D) and J20/VLW (E and F) mice. **(G and H)** Representation of the percentage of change in the percentage of PV-positive cells contacted by GABAergic SH fibers, and in the complexity of GABAergic SH contacts, comparing the four experimental groups. *n* = 4 animals per group, 3 sections per animal. Abbreviations: PV, Parvalbumin; so, *stratum oriens*; sp, *stratum pyramidale*; sr, *stratum radiatum*. Scale bar: 150 µm (A and B), 10 µm (C-F).

To determine whether the improvement in the GABAergic SH innervation in J20/VLW animals is due to a permanent recovery or a delay in the impairment of the GABAergic SH pathway, we analyzed 12 mo mice. Neither a reduction in the percentage of GAD- and PV-positive neurons contacted by GABAergic SH fibers nor in the number of synaptic boutons per GAD- and PV-positive cell was observed in 12 mo J20/VLW mice, compared to age-matched WT animals (Fig. 8).

**Figure 8.**
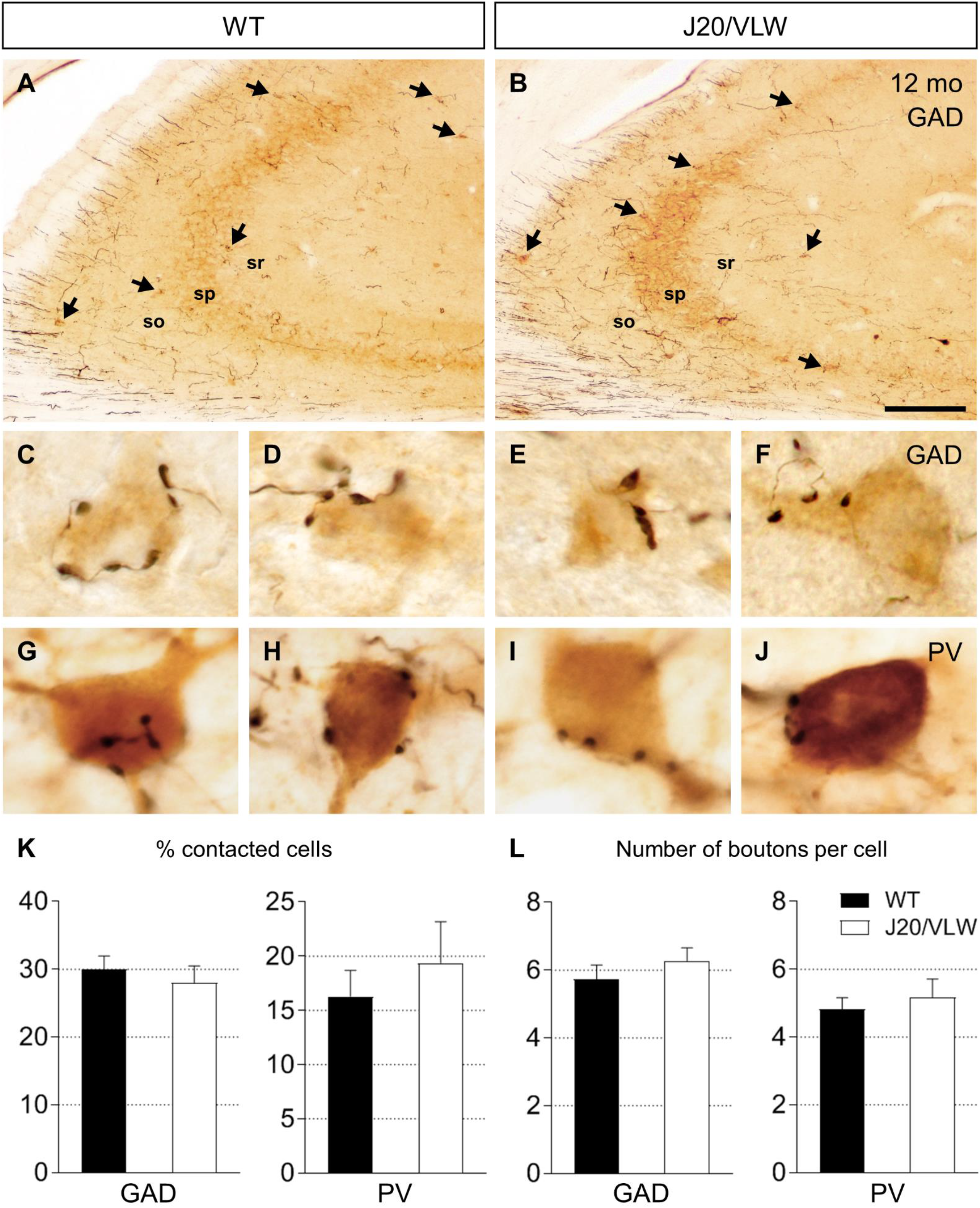
No alterations in GABAergic SH innervation are observed in 12 mo J20/VLW mice. Double immunodetection of GABAergic SH fibers and GAD- or PV-positive cells in hippocampal sections from 12 mo WT and J20/VLW mice. **(A and B)** GABAergic SH fibers (black) contacting GAD-positive cells (brown, arrows) in the CA3 region of WT (A) and J20/VLW (B) mice. **(C-F)** GABAergic SH baskets forming synaptic boutons on the soma of GAD-positive cells in WT (C and D) and J20/VLW (E and F) mice. **(G-J)** GABAergic SH baskets forming synaptic boutons on the soma of PV-positive cells in WT (G and H) and J20/VLW (I and J) mice. **(K and L)** Quantification of the percentage of GAD- and PV-positive cells contacted by GABAergic SH fibers (K), and the complexity of GABAergic SH contacts (L), in WT and J20/VLW mice. For (K) and (L): Student’s *t*-test. *n* = 4–5 animals per group, 3 sections per animal. Error bars represent SEM. Abbreviations: GAD, glutamic acid decarboxylase; PV, Parvalbumin; so, *stratum oriens*; sp, *stratum pyramidale*; sr, *stratum radiatum*. Scale bar: 150 µm (A and B), 10 µm (C-J).

These data suggest that the maintenance of correct GABAergic SH innervation in J20/VLW mice may contribute to the preservation of cognitive and physiological functions in this double transgenic mouse model. Our findings also suggest that the presence of Tau with a specific phosphorylation pattern, together with Aβ accumulation, in J20/VLW mice might confer neuroprotection against the GABAergic SH denervation associated to the individual Aβ and P-Tau pathologies.

To assess the GABAergic cell population in J20/VLW animals, we next analyzed the density and distribution of hippocampal and septal GABAergic neurons in 8 and 12 mo mice by immunodetection. The distribution and density of hippocampal GABAergic neurons (GAD-positive cells) in J20/VLW mice were similar to those of WT mice. GAD-positive cells were located throughout distinct layers and areas in the hippocampus (Supplementary Fig. 3A-D). We then studied the PV-positive subtype of interneurons. No alterations in either the distribution or density of PV-positive cells were detected in the J20/VLW hippocampus (Supplementary Fig. 3E-H).

Next, we analyzed the GABAergic SH neurons in the septal region by PV immunodetection. Our results indicated the preservation of both the distribution and density of PV-positive cells in the J20/VLW MSDB complex, compared to age-matched WT animals (Supplementary Fig. 4). We conclude that J20/VLW mice do not show altered distribution or loss of septal and hippocampal GABAergic neurons.

## DISCUSSION

Growing evidence indicates that Tau and Aβ have opposing effects on neuronal excitability and circuit activity (Angulo et al., 2017; Busche et al., 2019). However, the coexistence of Tau- and amyloid-related pathologies has been proposed to act synergistically to impair the function of neural circuits, and recent studies suggest that Tau has a dominating effect over Aβ (Angulo et al., 2017; Busche et al., 2019), which contrasts with previous findings (Ittner et al., 2010; Roberson et al., 2007). Here we show that the major alterations in theta and gamma rhythms and cognitive deficits observed in single transgenic J20 and VLW animals are not evident in J20/VLW mice. Our subsequent analyses suggest that the simultaneous presence of Tau phosphorylated at specific residues in hippocampal interneurons and of Aβ accumulation preserves hippocampal function in the double transgenic animals by maintaining a functional GABAergic SH pathway.

While Aβ may cause Tau phosphorylation and Tau may increase Aβ toxicity, there are conflicting lines of evidence as to whether Tau leads to an increase in amyloid deposition. Certain APP/Tau mice overexpressing hAPP^Sw^ together with hTau P301L (JNPL3/Tg2576 and APP23/B6P301L) show no differences in Aβ plaque load compared to single transgenic hAPP^Sw^ mice (Bolmont et al., 2007; Lewis et al., 2001). However, 16 mo Tg2576/VLW mice displayed enhanced amyloid deposition (Ribé et al., 2005). Our data demonstrate that no changes in Aβ deposition are present in J20/VLW mice compared to J20 animals. Previous studies demonstrated that Aβ increases Tau phosphorylation (Götz et al., 2001; Nisbet et al., 2015; Pérez et al., 2005). No changes in the levels of Tau phosphorylation at either residue Thr231 or residue Thr205 were observed in the present study in hippocampal pyramidal neurons when comparing VLW and J20/VLW mice, or at pSer262 Tau, which is absent in pyramidal neurons in both animal models. In addition, P-Tau mislocalization to the somatodendritic compartment of pyramidal neurons described previously in VLW mice (Dávila-Bouziguet et al., 2019; Lim et al., 2001; Soler et al., 2017) also occurred in the J20/VLW hippocampus. Thus, we conclude that the cognitive preservation observed in J20/VLW double transgenic animals is not due to a reduction in Aβ plaque load or to changes in the levels of P-Tau in pyramidal neurons.

In addition to the presence of P-Tau in pyramidal neurons (Götz et al., 1995; Rossi et al., 2020), hippocampal interneurons accumulate P-Tau in their soma in control and pathological conditions (Dávila-Bouziguet et al., 2019). As previously described in hAPP^Sw^/VLW mice (Pérez et al., 2005), our data indicate that Aβ presence enhanced Tau microtubule-binding domain phosphorylation at non-proline directed phosphorylation (NPDP) sites such as residue Ser262 in J20/VLW interneurons in comparison to VLW mice. These findings reveal that Aβ facilitates Tau phosphorylation at NPDP sites specifically in GABAergic neurons. It has been described that phosphorylation of Tau at residue Ser262 induces its detachment from microtubules (Ando et al., 2016). Moreover, it has been reported that Aβ induction of pThr231 Tau is dependent on pSer262 (Ando et al., 2016). Thus, the clear upward trend in the density of pThr231 Tau-positive interneurons in J20/VLW mice may result from the increased number of pSer262 Tau-positive interneurons in these animals. Taken together, these data suggest that, in the J20/VLW hippocampus, the increase in Tau phosphorylated at both residue Ser262 and residue Thr231 in GABAergic neurons facilitates the somatic localization of Tau, thereby favoring a novel function of this protein in the soma of hippocampal interneurons in this mouse model.

We previously described that Aβ accumulation in J20 mice and P-Tau presence in VLW animals induce abnormal GABAergic SH innervation in these two animal models (Rubio et al., 2012; Soler et al., 2017). Our results show that J20/VLW mice present neither altered distribution nor loss of septal or hippocampal GABAergic neurons, suggesting that the presence of P-Tau together with Aβ accumulation in J20/VLW mice protects against the GABAergic SH denervation associated with Aβ and P-Tau separately. Our findings also confirm a permanent effect since no alterations were observed in either 8 mo or 12 mo J20/VLW animals. No changes in Aβ plaque load or P-Tau accumulation and mislocalization in pyramidal neurons occurred in J20/VLW animals compared to J20 or VLW mice, respectively. Our data suggest that the maintenance of correct GABAergic SH innervation could be due to the specific pattern of Tau phosphorylation in hippocampal interneurons of J20/VLW mice.

Tau may participate in the dynamic regulation of GABA_A_ receptor trafficking at inhibitory synapses through the scaffolding protein Gephyrin, which is directly linked to the cytoskeleton (Essrich et al., 1998). By regulating GABA_A_ receptor clustering, Gephyrin controls GABAergic synaptic activity and, therefore, inhibitory transmission (Maric et al., 2017). In addition, it has been described that Glycogen Synthase Kinase 3β (GSK3β), a major Tau kinase activated by Aβ, regulates GABAergic synapse formation via the phosphorylation of Gephyrin (Tyagarajan et al., 2011; Tyagarajan and Fritschy, 2014).

Further research is required to gain a full understanding of the molecular mechanisms by which a distinct pattern of Tau phosphorylation modulates the function of GABAergic neurons. However, one hypothesis is that GSK3β activation by Aβ induces both an increase in P-Tau in the soma of GABAergic interneurons and the phosphorylation of Gephyrin, thereby contributing to GABA_A_ receptor clustering, thus preserving the GABAergic SH synaptic contacts on hippocampal interneurons and, therefore, stabilizing inhibitory synaptic activity.

Our previous data indicated that impaired GABAergic SH innervation in J20 mice correlates with altered patterns of neuronal hippocampal activity and with internal processes related to operant rewards. Spectral analysis showed a clear decrease in the spectral power of theta and gamma bands in J20 mice, compared to age-matched WT animals (Rubio et al., 2012; Vega-Flores et al., 2014). Furthermore, VLW animals overexpressing mutant hTau display hyperexcitability in the absence of Aβ, along with alterations in the GABAergic SH pathway (García-Cabrero et al., 2013; Soler et al., 2017). Here we demonstrate a considerable reduction in theta spectral power (33–42 %) in VLW animals. In contrast, J20/VLW mice showed only a slight decrease in the spectral power of theta (8–22 %) and gamma (33 %) bands. Aβ species cause synaptic loss and dysfunction in both glutamatergic and GABAergic synapses (Palop and Mucke, 2010; Rubio et al., 2012). It has been proposed that the GABAergic SH pathway regulates oscillatory activity, particularly through the recruitment of hippocampal interneurons. The main targets of the GABAergic SH fibers are the axo-axonic and basket PV-positive neurons, which control the firing of a large number of pyramidal neurons, hence leading to the generation of oscillatory activities in the range of the theta and gamma frequencies. Here we describe only minor alterations in the spectral power of theta and gamma bands in J20/VLW mice compared to J20 animals. Overall, our data indicate that the major alterations in theta and gamma rhythms observed in single transgenic J20 and VLW animals are only minor in J20/VLW mice, pointing to a correlation between preserved GABAergic SH innervation and proper hippocampal rhythmic activity.

The present study explores the relevance of correct GABAergic SH innervation and electrophysiological preservation for the cognitive state of J20/VLW animals. Our results indicate that the simultaneous presence of Aβ and P-Tau reverses the cognitive impairments observed in J20 and VLW mice. As described in AD and some AD animal models (Ambrad Giovannetti and Fuhrmann, 2019; Palop and Mucke, 2016; Verret et al., 2012), J20 and VLW animals display an imbalance between excitatory and inhibitory circuits associated with hyperexcitability and cognitive deficits. In contrast, double transgenic J20/VLW animals showed no major alterations in theta and gamma hippocampal rhythms, thereby suggesting a proper excitation/inhibition balance, probably modulated by GABAergic SH fibers, and, therefore, by correct hippocampal GABAergic function. Our results reveal no cognitive deficits in J20/VLW animals and point to a correlation between proper GABAergic synaptic function and cognition.

Taken together, our results suggest that the differential Tau phosphorylation pattern in hippocampal interneurons of J20/VLW mice protects against the loss of GABAergic SH innervation, thereby preventing alterations in LFPs and, subsequently, hindering cognitive deficits. These data support a new role of P-Tau in the maintenance of the GABAergic SH network and hippocampal GABAergic activity and indicate the potential of P-Tau regulation in GABAergic neurons as a therapeutic target in AD.

Recently, several research groups have described the existence of individuals presenting Aβ plaques and P-Tau tangles in the absence of cognitive impairment (Bjorklund et al., 2012; Briley et al., 2016; Erten-Lyons et al., 2009; Iacono et al., 2008; Singh et al., 2020; Zolochevska et al., 2018). These NDAN subjects are thought to have intrinsic mechanisms conferring protection against AD-associated dementia.

The understanding of the mechanisms underlying cognitive neuroprotection in the presence of both Aβ and P-Tau pathological traits could have a major impact in the design of therapeutic strategies aimed at preventing cognitive decline in AD patients. A major factor proposed as responsible for this protection is the preservation of the synaptic machinery that normally degenerates in AD (Bjorklund et al., 2012; Singh et al., 2020; Zolochevska et al., 2018). In this regard, we propose that the simultaneous presence of Aβ and Tau with a specific phosphorylation pattern in J20/VLW mice may confer protection against the synaptotoxic effects of pathogenic oligomers, as seen in NDAN individuals, or may trigger a differential pattern of gene expression that protects synaptic structure and function. Our results also point to the preservation of the GABAergic network as a critical factor underlying the recovery of cognitive and physiological deficits in J20/VLW mice. It is worth noting that the studies on NDAN subjects focus on glutamatergic synapses (Bjorklund et al., 2012; Singh et al., 2020; Zolochevska et al., 2018). Therefore, it would be of interest to analyze the state of GABAergic synapses in NDAN individuals to shed light on potential commonalities shared with J20/VLW mice.

Taken together, our results suggest that the differential Tau phosphorylation pattern in hippocampal interneurons of J20/VLW mice could protect against the loss of GABAergic SH innervation, thereby preventing alterations in LFPs and, subsequently, hindering cognitive deficits. These data support a new role of P-Tau in the maintenance of the GABAergic SH network and hippocampal GABAergic activity and indicate the potential of P-Tau regulation in GABAergic neurons as a therapeutic target in AD. Finally, we propose the double transgenic mouse line generated herein as a suitable animal model to understand the cognitive preservation in NDAN subjects and to open up new therapeutic strategies to treat AD-associated dementia.

## DISCLOSURE STATEMENT

The authors have no actual or potential conflicts of interest to disclose.

## ACKNOWLEDGEMENTS

The authors thank the personnel of the Advanced Optical Microscopy Facility at the Scientific and Technological Centers of the University of Barcelona for support, Alba Vílchez-Acosta for technical help, and the personnel of the Histopathology Facility of the Institute for Research in Biomedicine for assistance.

This work was supported by funds from the Ministry of Economy and Competitiveness (SAF2016-76340-R) and the Ministry of Science, Innovation and Universities (PID2019-106764RB-C21/AEI/10.13039/501100011033) to E.S., from the Ministry of Economy, Industry and Competitiveness (BFU2017-82375-R) to A.G. and J.M.D.-G; by the María de Maeztu Unit of Excellence awarded to the Institute of Neurosciences; and by an FPU grant from the Ministry of Education, Culture and Sport (FPU2016-07395) awarded to E.D.-B.

---

**Supplementary Figure 1.**
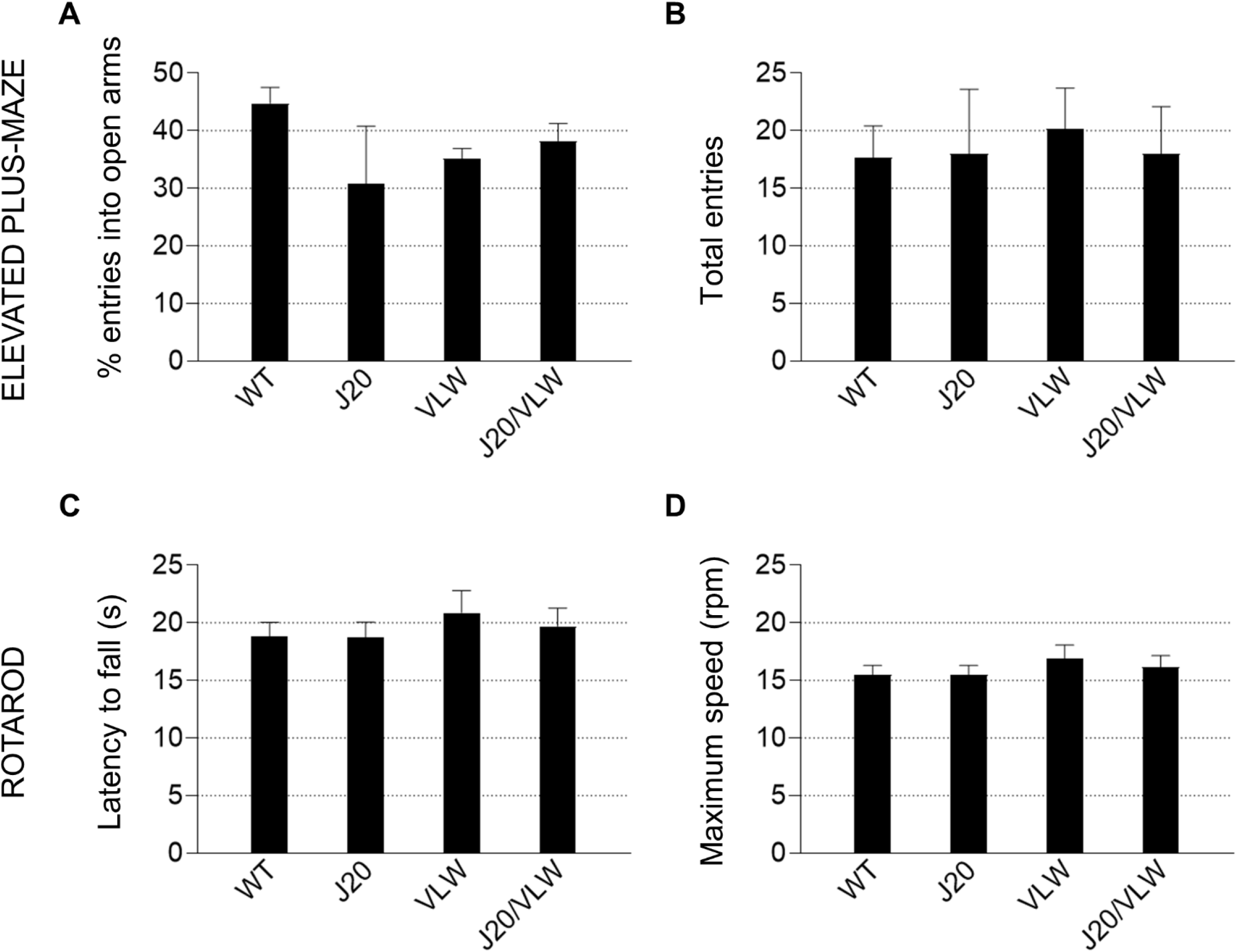
No major alterations in anxiety or motor coordination are present in J20/VLW mice. **(A and B)** Anxiety-like behavior was assessed using the elevated plus-maze and no significant differences were observed in either the percentage of entries into the open arms (A) or the total entries into closed and open arms (B) between the four experimental groups. **(C and D)** Motor coordination was measured using the accelerating rotarod and no alterations were observed in either the latency to fall (C) or the maximum speed (D) between the four experimental groups. For (A-D): one-way ANOVA. All behavioral tests were performed on 8 mo WT (*n* = 9), J20 (*n* = 6), VLW (*n* = 7), and J20/VLW (*n* = 5) mice. Error bars represent SEM.

**Supplementary Figure 2.**
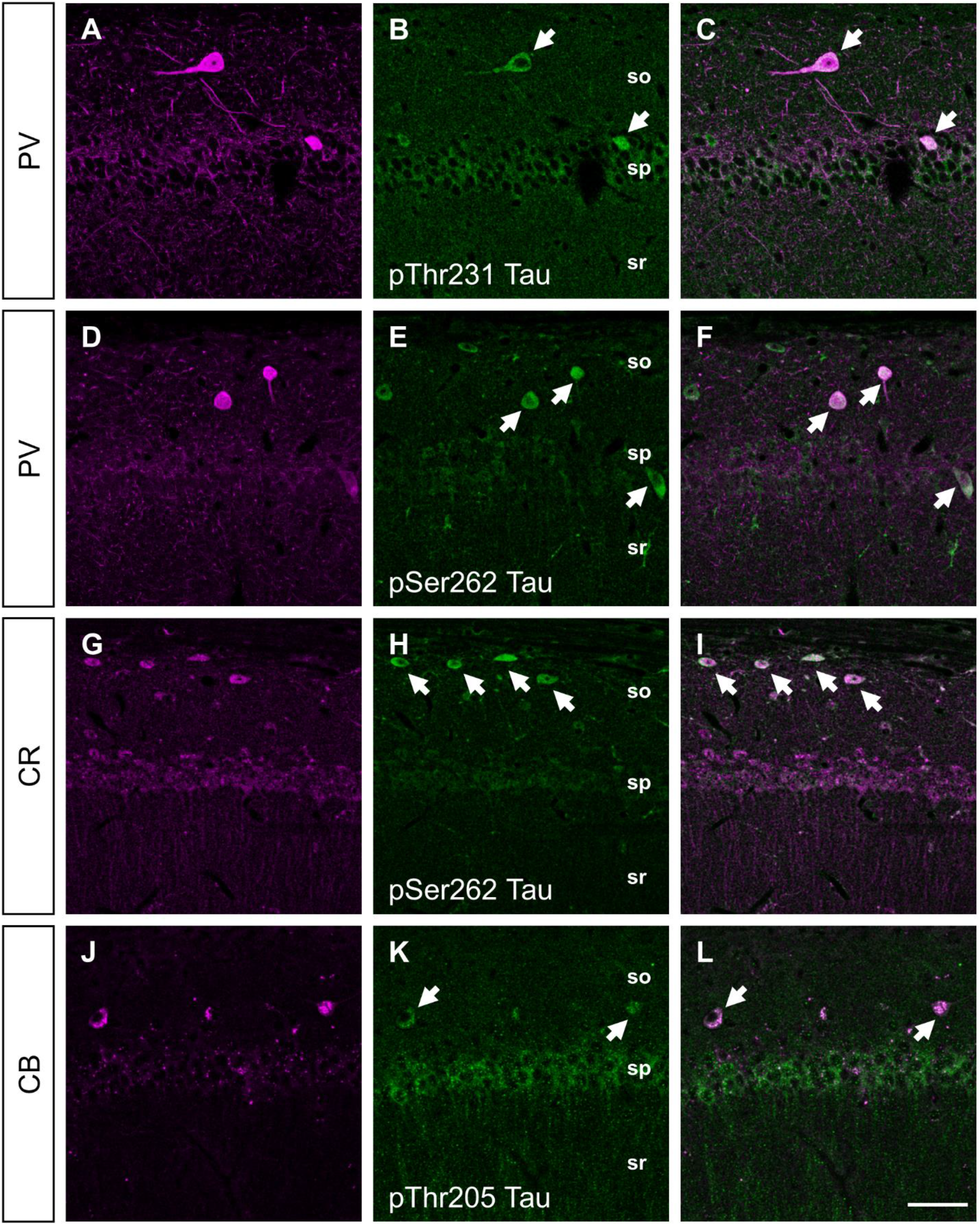
pThr231 Tau accumulates in PV-positive interneurons, and pThr205 and pSer262 Tau in PV-, CR-, and CB-positive interneurons in J20/VLW mice. Double immunofluorescent detection of pThr231, pThr205, or pSer262 and interneuron markers PV, CR, or CB in hippocampal sections from 8 mo J20/VLW mice. **(A-C)** PV-positive hippocampal interneurons of J20/VLW mice (magenta, A) accumulate Tau phosphorylated at residue Thr231 (green, B). Colocalization of pThr231 Tau with PV-positive interneurons (arrows in C). **(D-I)** PV- and CR-positive hippocampal interneurons of J20/VLW mice (magenta, D and G) accumulate Tau phosphorylated at residue Ser262 (green, E and H). Colocalization of pSer262 Tau with PV-(arrows in F) and CR-positive interneurons (arrows in I). (J-L) CB-positive hippocampal interneurons of J20/VLW mice (magenta, J) accumulate Tau phosphorylated at residue Thr205 (green, K). Colocalization of pThr205 Tau with CB-positive interneurons (arrows in L). Abbreviations: CB, Calbindin; CR, Calretinin; PV, Parvalbumin; so, *stratum oriens*; sp, *stratum pyramidale*; sr, *stratum radiatum*. Scale bar: 50 µm.

**Supplementary Figure 3.**
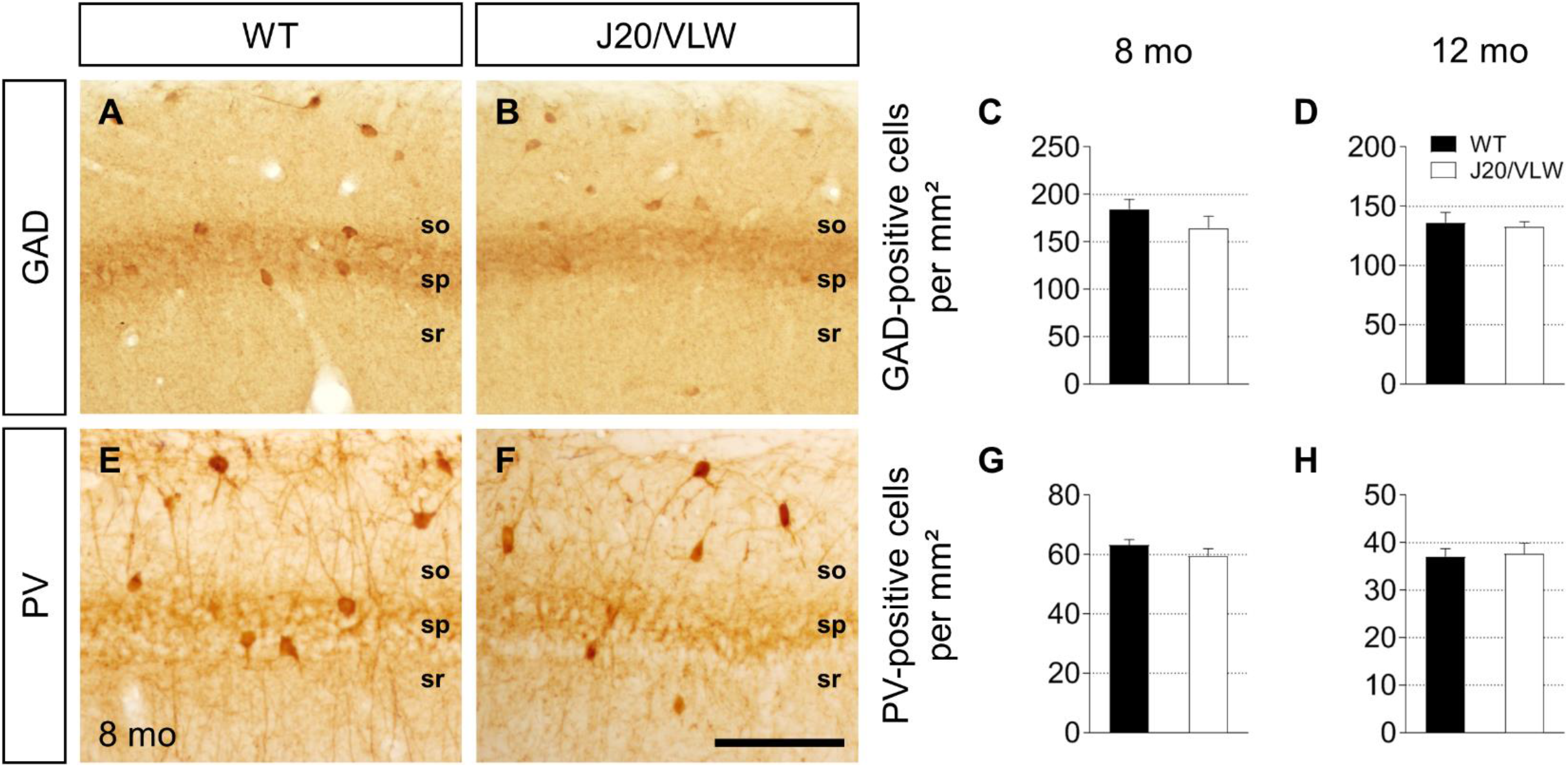
No loss of hippocampal GABAergic interneurons is observed in J20/VLW mice. Immunodetection and cell density quantification of the total GABAergic interneuron population (GAD-positive cells) and the PV-positive interneuron subpopulation in hippocampal sections from 8 mo and 12 mo WT and J20/VLW mice. **(A and B, E and F)** GAD-(A and B) and PV-positive cells (E and F) in the CA1 region of the hippocampus of 8 mo WT (A and E) and J20/VLW (B and F) mice. **(C and D)** Density quantification of GAD-positive cells in the hippocampus of 8 mo (C) and 12 mo (D) WT and J20/VLW mice. **(G and H)** Density quantification of PV-positive cells in the hippocampus of 8 mo (G) and 12 mo (H) WT and J20/VLW mice. For (C), (D), (G), and (H): Student’s *t*-test. *n* = 4–5 animals per group, 3 sections per animal. Error bars represent SEM. Abbreviations: GAD, glutamic acid decarboxylase; PV, Parvalbumin; so, *stratum oriens*; sp, *stratum pyramidale*; sr, *stratum radiatum*. Scale bar: 100 µm.

**Supplementary Figure 4.**
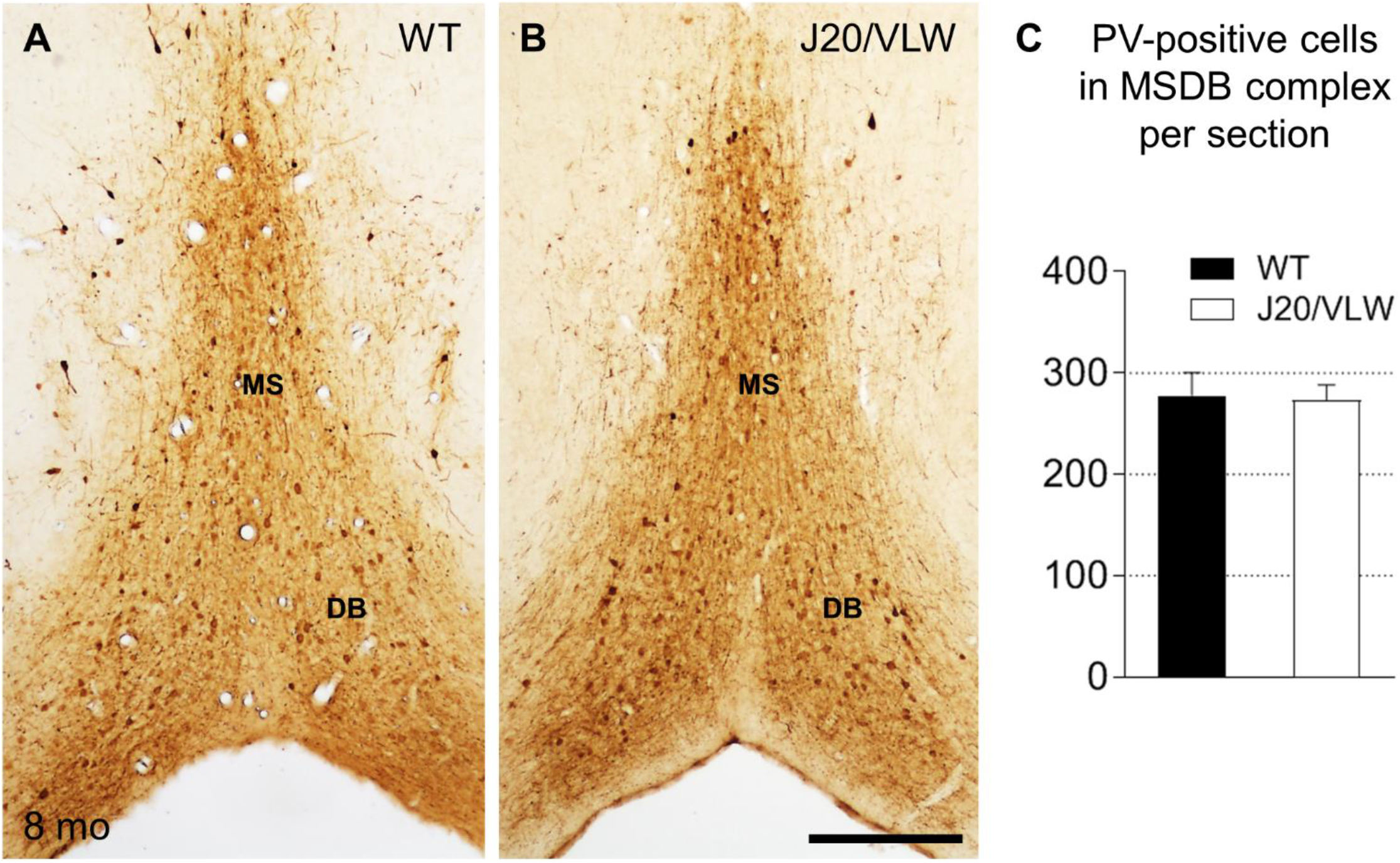
No loss of GABAergic SH neurons is observed in J20/VLW mice. Immunodetection of GABAergic SH neurons with PV antibody in septal sections from 8 mo WT and J20/VLW mice. **(A and B)** GABAergic SH cells, which express PV, in the MSDB complex of WT (A) and J20/VLW mice (B). **(C)** Density quantification of PV-positive cells in the MSDB complex of WT and J20/VLW mice. For (C): Student’s *t*-test. *n* = 4 animals per group, 4 sections per animal. Error bars represent SEM. Abbreviations: DB, nucleus of the diagonal band of Broca; MS, medial septal nucleus; PV, Parvalbumin. Scale bar: 300 µm.

## Notes

### Competing Interest Statement

The authors have declared no competing interest.

